# Identification of Maternal-Effect Genes in Zebrafish using Maternal Crispants

**DOI:** 10.1101/2021.02.19.432057

**Authors:** Cara E. Moravec, Gabriella C. Voit, Jarred Otterlee, Francisco Pelegri

**Affiliations:** Laboratory of Genetics, University of Wisconsin-Madison, Wisconsin

**Author notes:** Correspondence to: Francisco Pelegri, Laboratory of Genetics, 2424 Genetics/Biotech, University of Wisconsin –Madison, 425-G Henry Mall, Madison WI 53706.

**Keywords:** Crispr, maternal-effect, genome editing, early development, zebrafish, *kpna7*, *fhdc3*

## Abstract

In animals, early development is dependent on a pool of maternal factors, both RNA and proteins, which are required for basic cellular process and cell differentiation until zygotic genome activation. The role of a majority of these maternally expressed factors in adult fertility and early development is not fully understood. By exploiting the biallelic editing ability of CRISPR-Cas9 and the benefits of the zebrafish model, we identify and characterize maternal-effect genes in a single generation, using a maternal crispant technique. We validated the ability to generate biallelic mutations in the germline by creating maternal crispants that phenocopied previously characterized maternal-effect genes: *motley/birc5b*, *tmi*/*prc1l*, and *aura*/*mid1ip1*. Additionally, by targeting maternally expressed genes of unknown function in zebrafish, we identified two new maternal-effect zebrafish genes, *kpna7* and *fhcd3*. The genetic identity of these maternal crispants was confirmed by sequencing haploid progeny from F0 females, which allowed the sequence analysis of newly induced lesions in the maternal germ line. Analysis of the induced lesions shows minimal genetic variation within a clutch, with an average of two edited alleles per clutch. These findings are consistent with biallelic editing events occurring in germ cells or their precursors of early CRISPR-Cas9-injected embryos, leading to maternal-effect phenotypes in the offspring. Our studies show that maternal crispants allow for the effective identification and primary characterization of maternal-effect genes in a single generation, facilitating the reverse genetics analysis of maternal factors that drive embryonic development.

## Introduction

Model organisms have allowed for a greater understanding of early development, adult homeostasis, and human diseases because they permit testing for gene function within a developmental and molecular context. The zebrafish, *Danio rerio,* is an attractive vertebrate model due to its large number of progeny, external fertilization, and rapid transparent development. These features readily allow the assessment of loss-of-function conditions in embryonic development. Genes present in zebrafish are highly conserved, with approximately 70% of genes predicted to contribute to human disease having an ortholog in zebrafish (Howe et al., 2013). These benefits have led to the zebrafish becoming a valuable genetic model.

Recently the functions of many genes in zebrafish have been determined through reverse genetic approaches through the induction of loss-of-function conditions. Knockdowns have been achieved with the use of morpholino-conjugated oligonucleotides, although there is an uncertain correlation between morpholino and knockout phenotypes (Kok et al., 2015). The establishment and advancement of gene editing technology, such as zinc-finger nucleases (ZFNs)(Doyon et al., 2008; Meng et al., 2008), transcription activator-like effector nucleases (TALENs)(Huang et al., 2011), and clusters of regularly interspaced short palindromic repeats (CRISPR)(Hwang et al., 2013), have led to the ability to create targeted genetic lesions in candidate genes. The innovation of genome editing with CRISPR-Cas9 in zebrafish has expanded the number of potential targets in the genome at a significantly reduced effort and cost, thus making it widely accessible (Gagnon et al., 2014; Shah et al., 2015; Varshney et al., 2015a; Varshney et al., 2015b). Previous work has shown that embryos injected with CRISPR-Cas9 have the potential to contain biallelic editing, which allows for the screening of embryonic phenotypes in somatic tissue of injected (F0) embryos, known as a “crispant” (Burger et al., 2016; Jao et al., 2013; Shah et al., 2015; Shankaran et al., 2017; Trubiroha et al., 2018; Wu et al., 2018).

Early embryogenesis is dependent on a pool of maternal products, such as RNAs, proteins, and other biomolecules, which drive developmental processes until the activation of the zygotic genome (ZGA). Depletion of these maternal products, when essential, results in embryonic lethality. In animal species, mutations in maternally-expressed genes disrupt early cellular processes, such as egg polarity and activation, pronuclear fusion, cytokinesis, cell adhesion, and axis induction (Abrams and Mullins, 2009; Dosch, 2015). Maternally-expressed genes can be robustly targeted with CRISPR-Cas9 (Campbell et al., 2015; He et al., 2018; Li-Villarreal et al., 2015). However, an extra generation is necessary to observe maternal-effect phenotypes, which requires additional resources.

Here, we use maternal crispants to promote biallelic germline editing to achieve phenotyping of maternally-expressed gene function in the next generation. We first validated this technique by using maternal crispants to phenocopy three known maternal-effect genes: *motley/birc5b*, *tmi*/*prc1l,* and *aura*/*mid1ip1l* (Eno et al., 2016; Nair et al., Submitted; Nair et al., 2013). By using the zebrafish genome assembly to identify stage-specific RNA sequencing databases to predict expression and multiple-copy gene families, we selected four candidates with a potential maternal-only gene function. These candidates were tested for maternal-effect phenotypes using maternal crispants. Out of these four candidate genes, we identified two new zebrafish maternal-effect genes, *kpna7* and *fhcd3*. Overall, these studies demonstrate the feasibility of using available databases coupled with a maternal crispant approach to identify and characterize, in a single generation, novel maternal-effect genes in zebrafish.

## Materials and Methods

### Animal husbandry

Stocks of wild-type AB or crispant fish were raised and maintained under standard conditions at 28.5°C (Brand et al., 2002). Embryos were collected by natural matings and allowed to develop in E3 embryonic medium (Brand et al., 2002), with staging according to age and morphological standards (Kimmel et al., 1995). All zebrafish housing and experiments were approved by the University of Wisconsin-Madison Institutional Animal Care and Use Committee.

### Guide RNA Design and Synthesis

The CRISPR design web tool, CHOPCHOP, was used to identify guide RNA sites with high activity and minimal off-target sites (Labun et al., 2016; Montague et al., 2014; Moravec and Pelegri, 2019). For each target gene, four guide RNAs were selected with the following criteria: predicted to have high cleaving activity, located in or near the first predicted protein domain (as identified by Ensembl (Aken et al., 2016)), and minimal to no overlapping regions between guide RNAs.

Guide RNA templates were produced by using an annealing/fill-in method as described (Gagnon et al., 2014). Briefly, guide RNA templates were generated with a gene-specific oligo containing a SP6 promoter sequence, a gene-specific target sequence, and an overlapping sequence complementary to the constant oligo (Excel file 1). Annealing/fill-in was performed using T4 DNA polymerase using 100 μM of each of the oligo fragments. Oligos for each of the four gene-specific guide RNAs were separately annealed to the constant oligo. After the T4 DNA polymerase fill in, the four guide RNA templates were pooled together and concentrated through purification by a DNA Clean and Concentrator kit (Zymo Research).

A pooled guide RNA cocktail was synthesized using the MEGAshortscriptTM T7 Transcription Kit using the concentrated pooled guide RNA template. After RNA synthesis, the guide RNA pool was purified using an ethanol/ ammonium acetate protocol. The concentration of the guide RNA pool was measured by Nanodrop, and its integrity was checked by gel electrophoresis. Single-use aliquots of guide RNAs were stored at −80°C.

### Guide RNA/ Cas9 injection

A mixture of the pooled guide RNA cocktail (400pg/μl total final concentration) and Cas9 protein (PNA Bio; 200 pg/μl final concentration) was injected into one-cell stage AB embryos. At 24 hours, a subset of the injected embryos was collected, and DNA was extracted as previously described (Meeker et al., 2007). Guide activity was assayed by amplifying 120-100 bp fragment across the target site via PCR (Excel file 1) and then resolved on a 2.5% agarose gel. If Cas9-mediated genetic edits were created in the somatic cells, a population of DNA that contains a variety of INDELS would generate a smear, allowing the confirm Cas9 activity (Moravec and Pelegri, 2019).

### Phenotypic analysis of maternal crispant embryos

Injected embryos were raised until sexual maturity, and then all F0 females were mated through natural crosses to AB males. F1 embryos were collected approximately every 15 minutes and were allowed to develop in 1X embryonic medium (E3) in 100-mm Petri dishes, with no more than 100 embryos per 100 mm-diameter dish to ensure proper medium oxygenation. Control synchronized wild-type embryos and phenotypically wild-type embryos present in the F1 clutches allowed to further determine time from fertilization. Development was monitored in live embryos every hour for the first 8-10 hours, at 24 hours post-fertilization and twice a day thereafter until day five post-fertilization. Maternal crispants were scored for possible gross morphological defects, such as defective cellular division, deviations from the wild-type body plan, developmental delays, and failure to develop a swim bladder. For documentation of maternal-effect phenotypes, embryos less than 24 hours old were left in their chorions and imaged on glass-bottom dishes containing 1X E3, and embryos older than 24 hours were dechorionated and imaged on glass-bottom dishes containing 3% methylcellulose. All live images were acquired using a Leica MZ FLIII microscope equipped with a Leica EC3 camera and Leica Acquire version 1.0 RC4 software.

### Immunofluorescence and Microscopy

Embryos were fixed in 4% paraformaldehyde in phosphate buffered saline (PBS) and immunolabeled as previously described (Lindeman and Pelegri, 2012), with antibody incubation overnight at 4°C. Primary antibodies were anti-β-catenin made in rabbit (1:1000, Sigma C2206), anti-β-catenin made in mouse (1:1000, Sigma C7207), and anti-γ-Tubulin made in mouse (1:2000, Sigma T6557). Secondary antibodies were anti-Rabbit IgG made in goat, Alexa 488 (Molecular Probes A-11008), anti-Rabbit IgG made in goat, Cy3 (Jackson Immuno Labs 111-165-144), anti-mouse made in rabbit, Alexa 488 (Molecular Probes A-11059), and anti-mouse IgM made in goat, Cy3 (Jackson Immuno Labs 115-165-062). To detect DNA, embryos were labeled with a 0.5 μg/ml solution of 4′,6-diamidino-2-phenylindole-dihydrochloride (DAPI) for 10 minutes, followed by 3 PBS washes. After labeling, embryos were deyolked and mounted flat with the animal pole upwards in 75% glycerol. Images were obtained using a Zeiss LSM780 confocal microscope and analyzed using ImageJ.

### Analysis of *fhdc3* maternal crispants

To examine the shape of the early *fhdc3* and wild-type embryos, traces of the perimeters of embryos at 1.25 and 2 hours post-fertilization were analyzed using ImageJ. The presence of an inflated swim bladder was scored at five days post-fertilization as a sign of embryonic viability. Statistical analysis was performed using Student’s T-test.

### Sequencing of maternal crispants to identify genetic lesions

Females that produced maternal-crispant embryos were used for in vitro fertilization (IVF) for haploid production, as previously described (Baars et al., 2016). Briefly, the night before the IVF procedure, single F0 adult females were placed into mating boxes together with single wild-type AB males, with a physical barrier separating males and females. The morning of the IVF procedure, the barrier was removed, allowing natural mating. Tanks were visually monitored, and the F0 females were separated immediately after the first signs of egg release. In the same morning, a sperm solution was prepared using wild-type testes placed in Hanks’ solution (corresponding to testes from one male per 100 μl of Hanks’) and macerated, with the sperm released in solution treated with UV-light for 90 seconds to inactivate its DNA. Separated F0 females were anesthetized with tricaine solution (2% buffered tricaine in 100 ml of conditioned fish water), and mature eggs were extruded by applying gentle pressure to the abdomen. The extruded eggs were fertilized by the addition of 100 μl of UV-treated sperm solution with gentle mixing, followed by water-activation through the addition of E3 embryonic medium (1 ml for the first minute and flooding of the plate after the first minute). The resulting maternal haploid embryos were allowed to develop for 6-8 hours, with the time period allowing DNA content to increase through DNA replication in the developing embryos. After 6-8 hours, DNA was extracted from embryos that displayed a maternal crispant phenotype as previously described (Meeker et al., 2007), except that we used half of the volume (50 μl of 20mM NaOH and 5 μl of Tris-HCL pH 8.0) compared to the original protocol in order to concentrate the DNA.

To analyze genetic lesions in maternal haploids, DNA fragments corresponding to the targeted genes were PCR amplified using genomic DNA extracts from single haploid embryos. The PCR primers were designed to include multiple guide sites in order to capture potential site-spanning insertions or deletions (INDELS; Excel file 1). Duplicate PCR reactions of each sample were carried out and the amplified products were pooled using a DNA Clean and Concentrator kit (Zymo Research). The resulting amplified fragments were analyzed via Sanger sequencing using both forward and reverse primers. Sequence results were analyzed using Sequencher.

## Results

### Workflow for germline crispant approach and maternal-effect phenotyping

A traditional workflow for generating a mutant zebrafish with CRISPR-Cas9 uses guide RNAs targeting a specific gene of interest, co-injected in F0 early embryos with Cas9 mRNA or Cas9 protein. Successful targeting generates stable lines for the identification of a recessive zygotic mutation in F3 embryos, a process that requires three generations (in zebrafish, approximately nine to twelve months) from the time of injection. The identification of a mutant in a maternal-effect gene requires an additional generation to assess the phenotype in the progeny from a germ line with the loss-of-function condition (Figure 1A).

**Figure 1:**
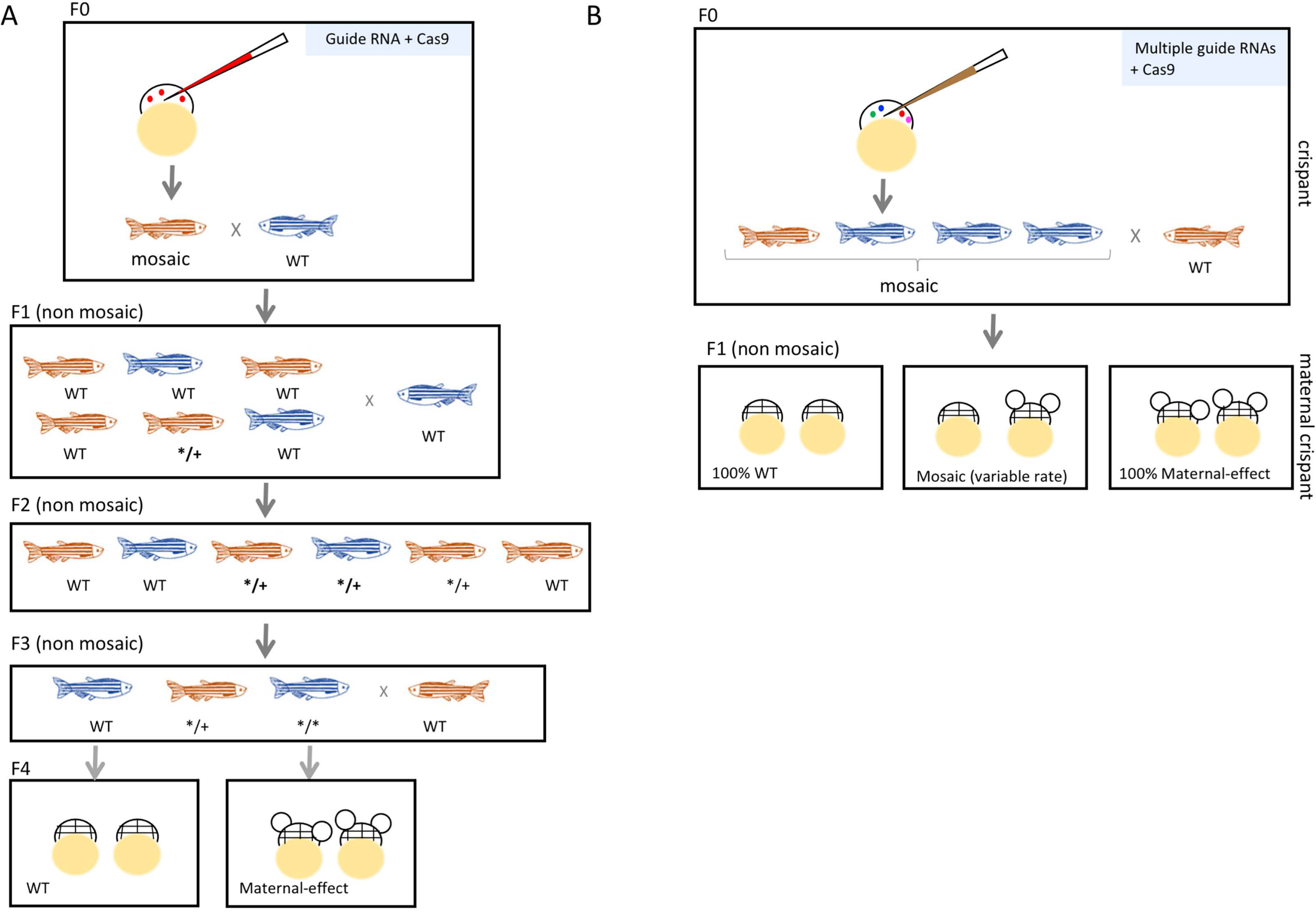
Maternal crispants can be used to efficiently identify maternal-effect genes in zebrafish a single generation. (A) Traditional workflow for CRISPR-Cas9 in zebrafish uses one guide RNA that is co-injected with Cas9 mRNA or protein into the one-cell embryo to target a specific gene of interest. Successful targeting will result in adults with a mosaic germ line. To isolate these alleles, non-mosaic F1 fish are generated by crossing the F0 CRISPR-injected fish with a wild-type fish, then screening for heterozygous carriers of INDELS. Outcrossing followed by incrossing results in female homozygous mutants for a maternal-effect gene, whose phenotypes can be assessed in their offspring. (B) A maternal crispant technique allows to obtain maternal-effect phenotypes in a single generation. This strategy multiplexes four guide RNAs to target a single gene, thus increasing the rate of biallelic editing. If this biallelic editing event occurs in the germline, the F1 offspring of F0 CRISPR-injected females have the potential to display a maternal-effect phenotype.

Recent work has shown that the biallelic editing ability of CRISPR-Cas9 allows for the screening of zygotic phenotypes in somatic cells of embryos in the F0 generation, also known as “crispants” (Burger et al., 2016; Jao et al., 2013; Shah et al., 2015; Shankaran et al., 2017; Trubiroha et al., 2018; Wu et al., 2018). Depending on the developmental process of interest, crispant embryos allow screening for the phenotype of targeted candidate genes in a few hours to a few days. In zebrafish, multiplexing guide RNAs to a single gene target has been shown to generate biallelic lesions with high efficiency (Wu et al., 2018). This increased efficiency should in principle allow the observation of maternal-effect phenotypes in the F1 generation in what we refer to as “maternal crispant” embryos. The generation of maternal crispants would allow testing the phenotype of mutations in novel maternally-expressed genes directly in the progeny of injected individuals.

For maternal crispants to occur, there must be biallelic editing events in cells fated to develop into the germline. To generate maternal crispants with high efficiency, we designed four guide RNAs for each target, which included known or candidate maternal-effect genes. Guide RNAs were typically designed to target the first conserved domain of the protein, as listed on Ensembl, to increase the probability that any INDELS, both in-frame and out-frame shifts, abolish protein function. To generate maternal crispants, a cocktail containing multiplexed guide RNAs targeting a single gene and Cas9 protein was injected into one-cell (F0) embryos. Once the injected F0 embryos reached adulthood, the resulting females (heretofore referred to as “F0 females”) were screened for maternal-effect phenotypes by scoring their offspring for developmental abnormalities. The resultant clutches from F0 females, presumably depending on the fraction of the germline containing biallelic lesions, exhibited a range of phenotypes, from 0 to 100% affected embryos (Figure 1B).

### Validation of the maternal crispant approach using known maternal-effect genes

Three known maternal-effect genes, *motley*/ *birc5b*, *tmi*/*prc1l*, and *aura*/ *mid1ip1l,* which when mutated result in defects in cytokinesis in the early embryonic cell divisions (Eno et al., 2016; Nair et al., Submitted; Nair et al., 2013), were chosen to validate a maternal crispant approach. Guide RNAs targeted the conserved regions for *birc5b* (BIR domain; Figure 2A); and *prc1l* (start of the MAP65/Ase/PRC1 domain; Figure 2B), and the single exon for *mid1ip1l* (Figure 2C). F0 females were mated to wild-type males, and the resulting F1 clutches were examined for phenotypes. In nearly all cases (9 of 10 F0 females), F1 clutches contained embryos with defects indistinguishable from those of embryos derived from females homozygous for the corresponding target gene (*birc5b*: 78-100%; *prc1l*; 27-47%; *mid1ip1l*: 0-60%) (Table 1). Two of the F0 females (for *birc5b*) produced 100% maternal crispant embryos, indicating that in these females most if not the entire germ line contained biallelic edits. The remaining F0 females produced a fraction of embryos exhibiting the target mutant phenotype as well as wild-type embryos, reflecting a mosaic germline. One F0 female (for *mid1ip1l*) did not yield maternal crispant embryos.

**Figure 2:**
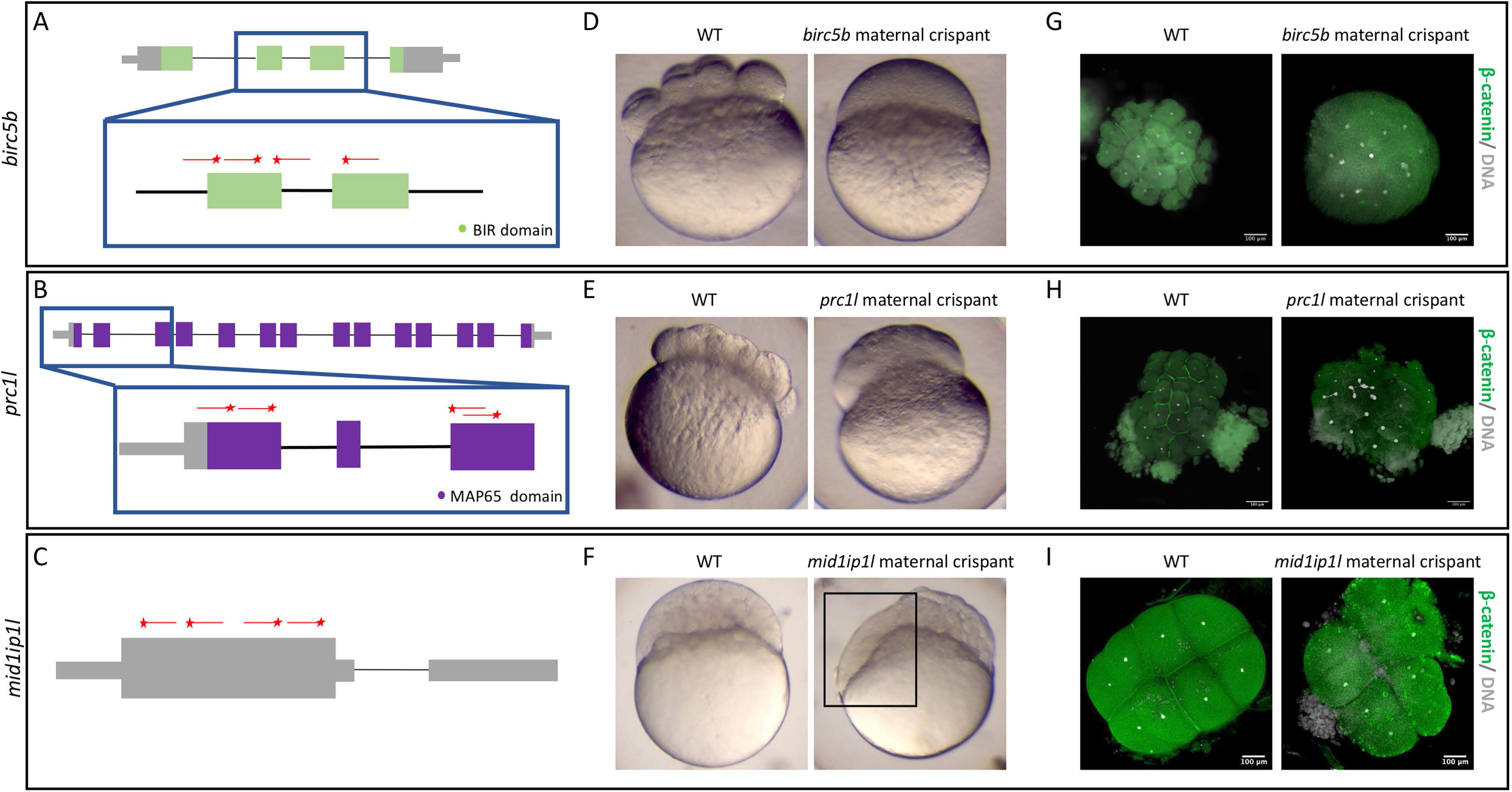
Identification of known maternal-effect genes by using maternal crispants. Known maternal-effect phenotypes resulting from the mutagenesis of three genes, *motley*/ *birc5b*(A,D,G)*, tmi/ pcr1l*(B,E,H), and *aura/ mid1ip1l* (C,F,I), were replicated using the maternal crispant technique. A-C) Gene structure diagrams of *birc5b* (A), *prc1l* (B), and *mid1ip1l* (C) showing the target sites of guide RNAs (red lines) and PAM sites (red stars). D-F) Representative comparisons of a live wild-type embryo with a live time-matched *birc5b* maternal crispant embryos of *birc5b* (D, 75 minutes post fertilization (mpf)), *prc1l* (E, 75mpf), and *midip1il* (F, 150 mpf), showing defects in cytokinesis identical to previous studies. *mid1ip1l* maternal crispant embryo shown exhibits a partially syncytial phenotype (box in F), as previously reported. G-I) Immunohistochemistry labeling of anti-β-catenin and DAPI of birc5b (G, 90 mpf), prc1l (H, 90 mpf), and midip1l (I, 75mpf) shows the lack of accumulation of β - catenin in mature furrows, confirming a failure in furrow formation in spite of DNA replication. *mid1ip1l* maternal crispants contain ectopic β-catenin-containing vesicles near the cortex, as previously reported. Magnifications are shown with scale bars, with some embryo size variation caused by methanol storage of embryos in (G,H).

**Table 1:** Fraction of embryos exhibiting a maternal crispant phenotype for targeted previously known maternal-effect genes, *birc5b*, *prc1l,* and *midlip1l*.

| Gene | Fish | % Phenotype | N |
| --- | --- | --- | --- |
| *birc5b* | 1 | 78 | 14 |
|  | 2 | 100 | 145 |
|  | 3 | 100 | 43 |
| *prc1L* | 1 | 27 | 48 |
|  | 2 | 47 | 82 |
|  | 3 | 47 | 91 |
| *mid1ip1l* | 1 | 33 | 51 |
|  | 2 | 60 | 82 |
|  | 3 | 12 | 56 |
|  | 4 | 0 | 30 |

Characterization of the maternal crispants confirmed cellular defects identical to those caused by loss of function of the corresponding target genes. Like embryos from *motley* homozygous females (Nair et al., 2013), *birc5b* maternal crispant embryos fail to create a visible furrow (Figure 2D) and in spite of normal DNA segregation fail to accumulate the cell adhesion component ß-catenin at furrow sites (Figure 2G). Similar to embryos from females homozygous for *tmi* mutations (Nair et al., Submitted), *prc1l* maternal crispant embryos showed incipient cleavage furrows that undergo regression (Figure 2E), exhibiting reduced β-catenin deposition at the furrow and aberrant chromosome segregation (Figure 2H). Corresponding to embryos from females homozygous for *aura* mutations (Eno et al., 2016), *mid1ipil* maternal crispants exhibit abnormal cell division and become partially syncytial (Figure 2F), with ß-catenin present in unreleased cortical granules and unavailable for recruitment to the furrow (Figure 2I).

In summary, for all of three targeted known maternal-effect genes, a maternal crispant approach was able to reliably replicate loss-of-function phenotypes.

### Identification of novel maternal-effect genes via maternal crispants

The ability to reliably identify maternal crispant embryos from F0 females for three tested known maternal-effect genes provided proof-of-principle for a maternal crispant approach. Maternal-specific genes typically have high transcript levels in the early embryo with a severe reduction in mRNA levels as the embryo undergoes zygotic genome activation (ZGA) (Aanes et al., 2011). In addition, a number of characterized maternal-effect mutations in zebrafish occur in genes that arose through gene duplication events, with the gene affected by the maternal-effect mutation expressed largely or exclusively maternally, and other copies, presumably the ancestral ones, retaining a more widespread function (Campbell et al., 2015; Eno et al., 2016; Ge et al., 2014; Nair et al., Submitted; Nair et al., 2013). This pattern is consistent with the functional divergence of duplicated genes (Sandve et al., 2018), which for some genes acquire a maternal-only function. To identify candidates to target using a maternal crispant approach, we used an RNA-seq database that provides a baseline transcriptional profile for multiple developmental stages of the zebrafish, from zygote to 5-days post-fertilization (White et al., 2017). We searched for uncharacterized genes that: 1) displayed an expected profile for a maternal-specific gene, being expressed in the oocyte and early embryo and degraded at ZGA, and 2) are gene duplicates where one or more additional copies are expressed maternally and zygotically, or zygotically only, reflecting a likely role for this second gene duplicate throughout development. These criteria are expected to enrich for maternally-expressed gene duplicates whose knock out would have no effect on zygotic development in F0 fish but could lead to maternal-effect phenotypes. Using these criteria, we selected four candidate maternal-effect genes: *kpna7*, *fhdc3*, *zgc:114123*, and *kcns3b*. These genes are predicted to carry out a diverse set of cellular functions in the zebrafish embryo, related to nuclear import (*kpna7*), regulation of actin dynamics (*fhdc3* and *zgc:114123),* and maintenance of cell membrane potential (*kcns3b).*

The application of the maternal crispant technique to these genes allowed for the identification of maternal-effect phenotypes for two of these genes, *kpna7* and *fhdc3*. Targeting of *zgc:114123* and *kcn3b* led to clutches from F0 injected females that contained only phenotypically wild-type embryos (Supplementary Table 1, Supplementary Figure 1), in spite of the production of multiple INDELS in F0 embryo somatic tissues 24 hours after injection (data not shown).

### Kpna7 is necessary for nuclear stability in the early embryo

In eukaryotic cells, large proteins are actively transported from the cytoplasm into the nucleus via a transport system mediated by a heterodimeric Importin α/Importin β complex that recognizes cargo containing a nuclear localization signal (NLS)(Conti et al., 1998; Görlich et al., 1994; Imamoto et al., 1995; Moroianu et al., 1995; Yasuhara et al., 2004).

Eukaryotes contain a varying number of Importin α genes, for example yeast has a single *kpna* gene, while mammals have seven *kpna* genes (Hu et al., 2010; Kelley et al., 2010; Tejomurtula et al., 2009). Among mammalian *kpna* genes, only *kpna7* shows expression in the oocyte and the early embryo and RNA degradation at ZGA (Figure 3A)(Boroviak et al., 2018; Hu et al., 2010; Tejomurtula et al., 2009; Wang et al., 2012). Similar to mammals, zebrafish also have seven *kpna* genes (Figure 3B), with *kpna7* exhibiting maternal-specific expression and transcript degradation at ZGA (Figure 3B). We targeted zebrafish *kpna7*, which has a canonical structure for this family of proteins composed of an N-terminal Importin-β binding domain and multiple armadillo repeats at the C-terminus (Figure 3C)(Conti et al., 1998; Görlich et al., 1994).

**Figure 3:**
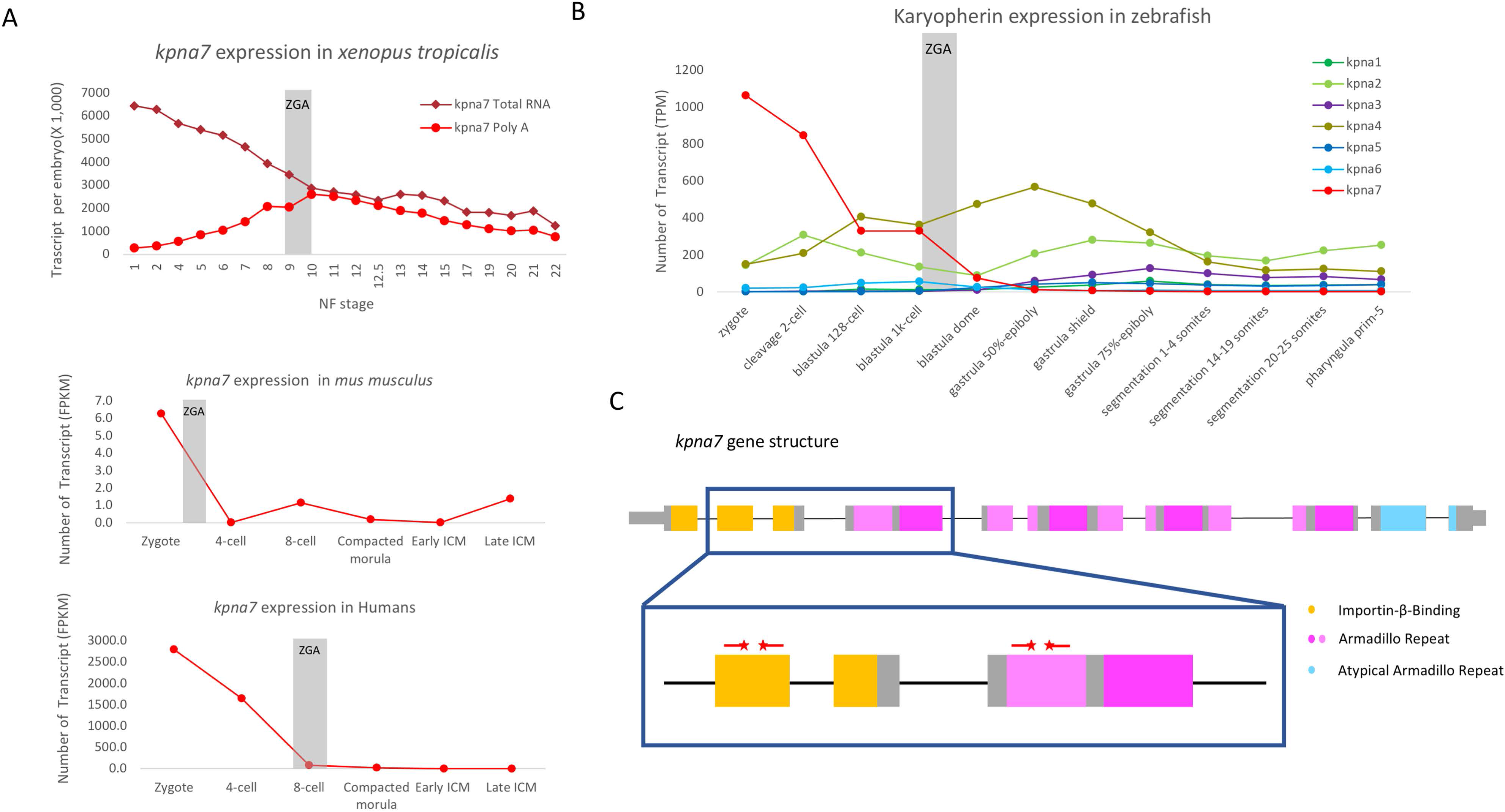
Gene structure and expression of *kpna7* during development. A) Expression levels of *kpna7* in other species display a similar pattern of the maternal expression of *kpna7* homologues. B) Expression levels of the *kpna* family throughout zebrafish development, from zygote to Prim-5 (24hpf) stage. The maternal-specific transcript, *kpna7*, is represented in red and gray bar marks the zygotic genome activation. C) A diagram of the gene structure of *kpna7*, showing the known motifs and target sites of guide RNAs (red lines) and PAM sites (red stars).

For the zebrafish *kpna7* maternal crispant, two guide RNAs were designed to target the Importin-β binding domain and two additional guide RNAs to target the first Armadillo repeat (Figure 3C). Clutches from two different *kpna7* F0 females exhibited a maternal phenotype in 100% of the embryos (Table 2). During the cleavage stages, the *kpna7* maternal crispant embryos were morphologically indistinguishable from wild-type embryos. However, as they undergo development, the majority of these embryos stalled at sphere stage, with only a small fraction of embryos initiating epiboly and none completing this process (Figure 4A). Labeling for β-catenin and DAPI at 6 hours post fertilization (hpf) shows that *kpna7* maternal crispant embryos exhibit an uneven distribution of nuclei, with a subset of cells appearing to entirely lack DNA (Figure 4B, Movie 1)(n=15) and others exhibiting nuclei of a range of sizes and shapes, in contrast to their even distribution and size in wild-type (Figure 4B, Movie 2)(n=16).

**Table 2:** Fraction of embryos exhibiting a maternal crispant phenotype for candidate genes identified using gene expression and phylogenetic criteria, *kpna7* and *fhdc3*.

| Gene | Fish | % Phenotype | N |
| --- | --- | --- | --- |
| *kpna7*<br>(stalled<br>development<br>at sphere ) | WT | 0 | 111 |
|  | 1 | 100 | 40 |
|  | 2 | 100 | 25 |
| *fhdc3*<br>(yolk in<br>blastodisc at<br>sphere) | WT | 1 | 133 |
|  | 1 | 74 | 43 |
|  | 2 | 100 | 51 |
|  | 3 | 97 | 118 |

**Figure 4:**
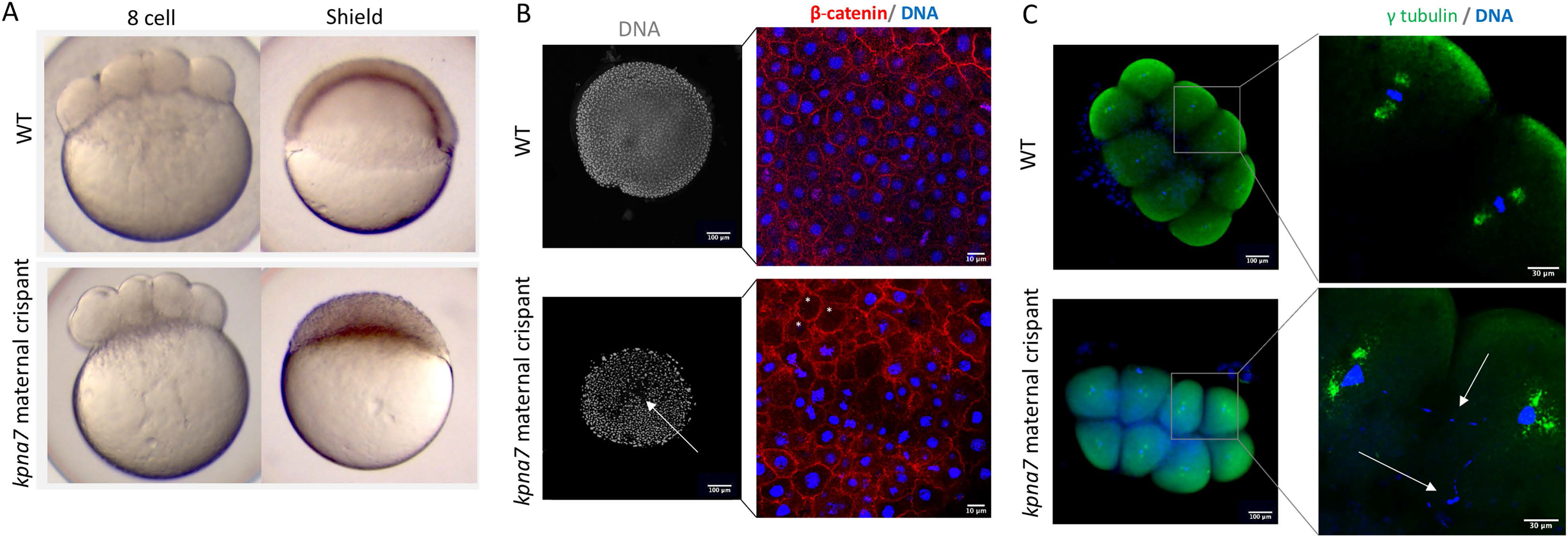
Kpna7 is necessary for nuclear segregation during early development. A) Live images of *kpna7* maternal crispant and wild-type controls at the 8-cell stage (75 mpf) and shield stage (6hpf). At the 8-cell stage, *kpna7* maternal crispants appear to divide normally, but they stall at the sphere stage and fail to undergo epiboly. B) Immunohistochemistry labeling of anti-β-catenin and DAPI at 6 hpf showing that the *kpna7* maternal crispant embryos exhibit nuclei of unequal sizes including a subset of cells that entirely lack nuclei (asterisks). C) Immunohistochemistry labeling of anti-γ-tubulin and DAPI at 75 mpf, showing that *kpna7* maternal crispant embryos display abnormal nuclear segregation during cell cleavage.

In contrast to *kpna7* knockdown cancer cell lines, which exhibit supernumerary centrosomes (Vuorinen et al., 2018), all cells in *kpna7* maternal crispants showed the normal centrosome pair component (Figure 4C; wild type: n=44 cells in 6 embryos; *kpna7* maternal crispants: n=39 cells in 7 embryos). However, *kpna7* maternal crispants exhibited defective DNA morphology in a majority of cells, reflected in nuclear material that failed to segregate to the spindle poles (Figure 4C; wild-type: 0/22 abnormal DNA segregation events in 6 embryos; *knpa7* maternal crispants: 17/21 abnormal DNA segregation events in 7 embryos). This defect, likely compounded through multiple rounds of cell division, presumably results in the uneven nuclear size and distribution observed at later stages. Although more studies will be needed to study its function, this work indicates that maternal-expressed zebrafish *kpna7* is necessary for DNA morphology and/or segregation in the early embryo.

### Fhdc3 is necessary for the yolk-blastodisc boundary function

Proteins of the Formin family are key regulators of cytoskeletal reorganization through the promotion of actin nucleation in the cell, with a well-established role in processes such as cytokinesis, cell polarization, and filopodia assembly (Breitsprecher and Goode, 2013; Goode and Eck, 2007; Hegsted et al., 2017). A subset of proteins in the Formin family are known as ‘inverted formins”, where the regulatory N-terminal region has been lost leaving the FH1 and FH2 domains at the N-terminal region of the protein (Breitsprecher and Goode, 2013; Hegsted et al., 2017; Higgs and Peterson, 2005). Inverted formins have been shown to be involved in the linking of F-actin and microtubules and to have a role in membrane trafficking and lumen formation (Breitsprecher and Goode, 2013; Hegsted et al., 2017; Higgs and Peterson, 2005; Young et al., 2008).

In zebrafish, the *fhdc* family is composed of 5 different genes, *fhdc1*, *fhdc2*, *fhdc3*, *fhdc4*, and *fhdc5*, with *fhdc3* as its sole member with maternal-only expression (Figure 5A, data not shown). To generate *fhdc3* maternal crispants, guide RNAs were designed with one guide RNA targeting the protein before the FH1 domain and the other three guide RNAs targeting the start of the FH2 domain (Figure 5B). Three *fhdc3* F0 females were screened, and all three generated clutches containing embryos exhibiting a consistent maternal-effect phenotype (Table 2). These embryos undergo normal cell cleavage during the first few cell cycles but, at approximately the 32-cell/64-cell stage, start to exhibit an aberrant constriction at the yolk-blastodisc boundary (black arrowhead) (Figure 6A). Immunolabeling of *fhdc3* maternal crispants with β-catenin and DAPI shows that at the 8-cell stage, the cellular layout (a 2×4 cell-matrix at this stage) is similar between the *fhdc3* maternal crispants and stage-matched wild-type controls. However, at the 64-cell stage, when wild-type embryos exhibit a regular cellular layout (organized largely as 2 tiers of blastomeres, each in a 4×8 cell-matrix), *fhdc3* maternal crispant embryos exhibit an unorganized cellular layout (Figure 6C). The blastoderm of early wild-type embryos exhibits an oblong shape (Kimmel et al., 1995), possibly caused by the coupling of blastomere division to spindle lengthening (Wühr et al., 2010; our own observations). Using a score of circularity (with a perfect circle having a score of 1) to broadly assess blastoderm shape, we found that, starting at the 64-cell stage, *fhdc3* maternal crispant embryos are significantly less oblong than wild-type (Figure 6D). Altogether our observations indicate that embryos lacking *fhdc3* function are unable to maintain the ordered cellular arrangement of the early wild-type blastoderm.

**Figure 5:**
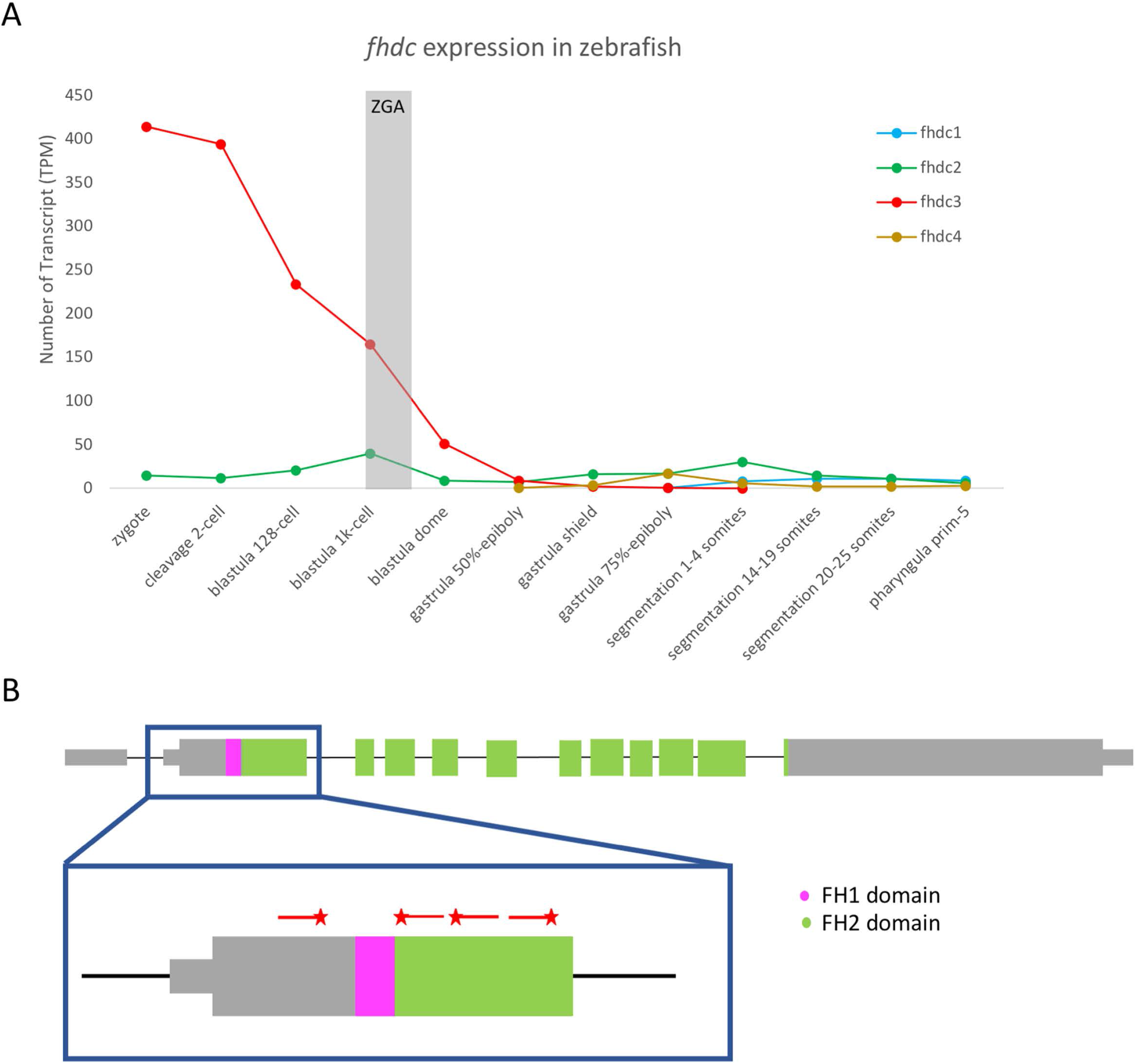
Maternal expression (A) and gene targeting (B) of *fhdc3* during development. Figure legend as in Fig. 3C. Expression data in early development in other systems is not yet known.

**Figure 6:**
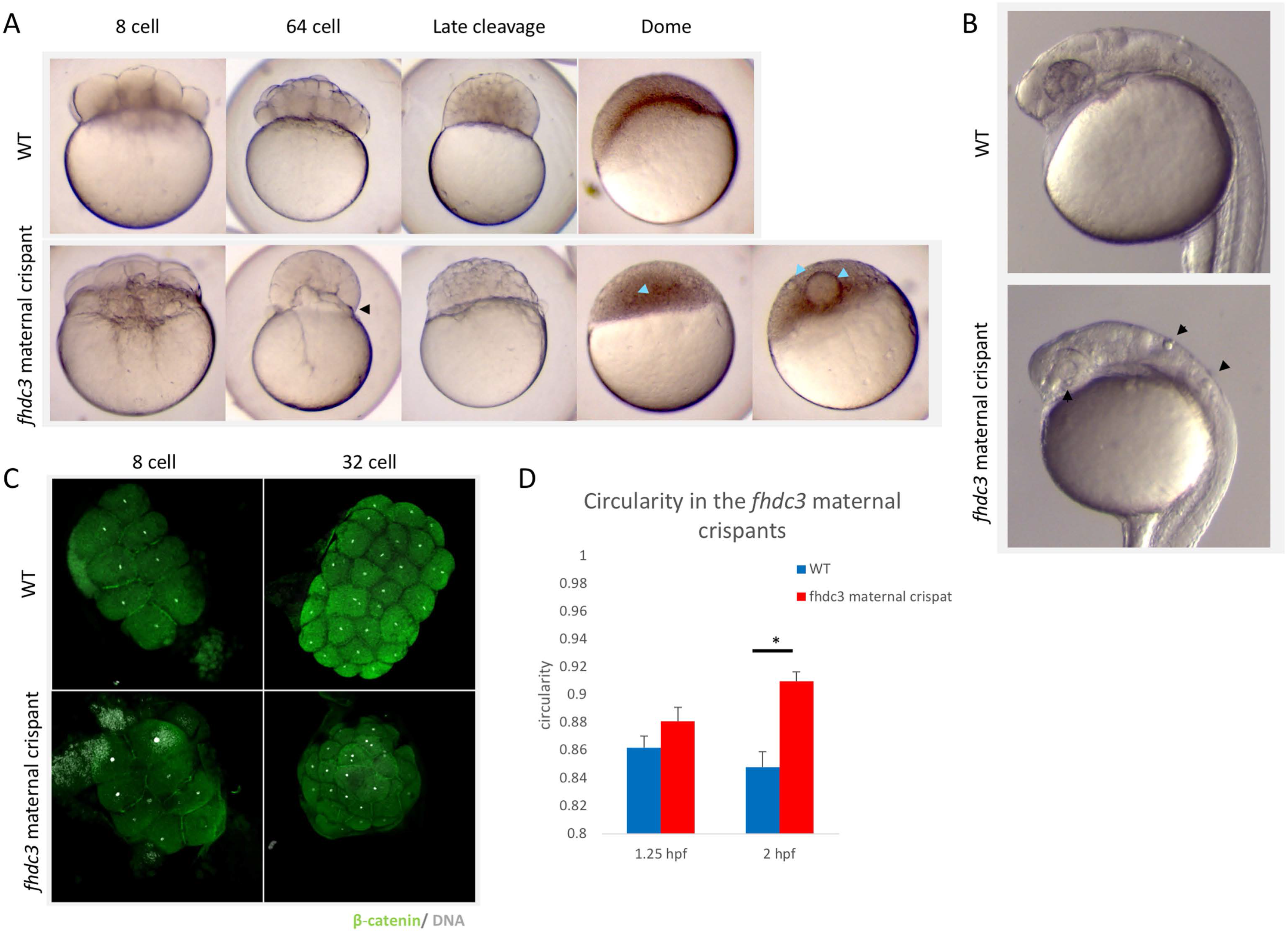
Fhdc3 is necessary to maintain the yolk:blastodisc boundary during early development. A) Live images comparing *fhdc3* maternal crispant embryos to time-matched wild-type controls during embryonic development. An abnormal constriction at the boundary between the blastodisc and yolk (black arrowhead) is observed during the early cleavage stages (see 64-cell). At later stages a normal degree of constriction at the yolk-blastodisc boundary is restored, but embryos exhibit ectopic yolk inclusions (light blue arrowheads). B) One day post-fertilization, *fhdc3* maternal crispant embryos do not exhibit gross morphological defects although still contain ectopic yolk inclusions (black arrowheads.) C) Immunolabeling with anti-β-catenin and DAPI at 8-cell and 64-cell stages, showing that the overall shape and cell-organization of the embryo is affected in the later *fhdc3* maternal crispant embryos. D) At the 8-cell stage, there is no significant difference in the shape of wild-type and *fhdc3* maternal crispant embryos (*fhdc3* maternal crispant score of circularity 0.881 (n=18); wild type 0.8615 (n=13); P=.0893) but the shape appears significantly different at the 64-cell stage (*fhdc3* maternal crispants: 0.90963 (n=8), wild type: 0.84757 (n=7); P=.0003). Error bars represent standard error.

In spite of earlier defects, by late-cleavage stages *fhdc3* maternal crispants no longer show an abnormal constriction of the yolk-blastoderm boundary phenotype, appearing instead at these stages morphologically relatively normal (Figure 6A). Nevertheless, at these stages *fhdc3* maternal crispant embryos contain ectopic yolk inclusions in varying numbers and sizes embedded in the blastodisc, presumably from a defective cytoplasm-yolk boundary at earlier stages (Figure 6A). Even with these inclusions, *fhdc3* maternal crispants are able to proceed through gastrulation without apparent gross morphological defects up to 5 days (Figure 6B, data not shown).

A closer examination of *fhdc3* maternal crispants at 24 hpf shows the continuing presence of ectopic yolk inclusions distributed in various regions of the embryo (Figure 6B; black arrowheads, data not shown), which persist in the embryo up to at least 5 days post-fertilization (data not shown). *fhdc3* maternal crispant embryos that contain inclusions are significantly less likely than wild-type controls to be viable at five days post-fertilization, as assessed by swim bladder inflation (wild type: 96%, n=72; *fhdc3* maternal crispants: 39%, n=109; P-value = 0.009).

Our observations indicate that *fhdc3* plays an essential role in regulating the boundary between the yolk and the blastodisc in the early embryo, required for the segregation of the yolk away from the blastodisc and eventually embryonic survival.

### Identification of genetic lesions in maternal crispants mutants

The multiple populations of alleles in traditional crispants, which typically occur in diploid cells and additionally involve a mixture of induced lesions, can lead to challenges in interpreting the sequences of targeted genes. To facilitate the analysis of lesions in maternal crispants, we took advantage of the ability to generate haploid embryos in zebrafish (Walker, 1998) and sequenced alleles present in single haploid maternal crispant embryos. This combined approach allowed for straightforward identification and characterization of genetic changes to the target sites via Sanger sequencing (Figure 7).

**Figure 7:**
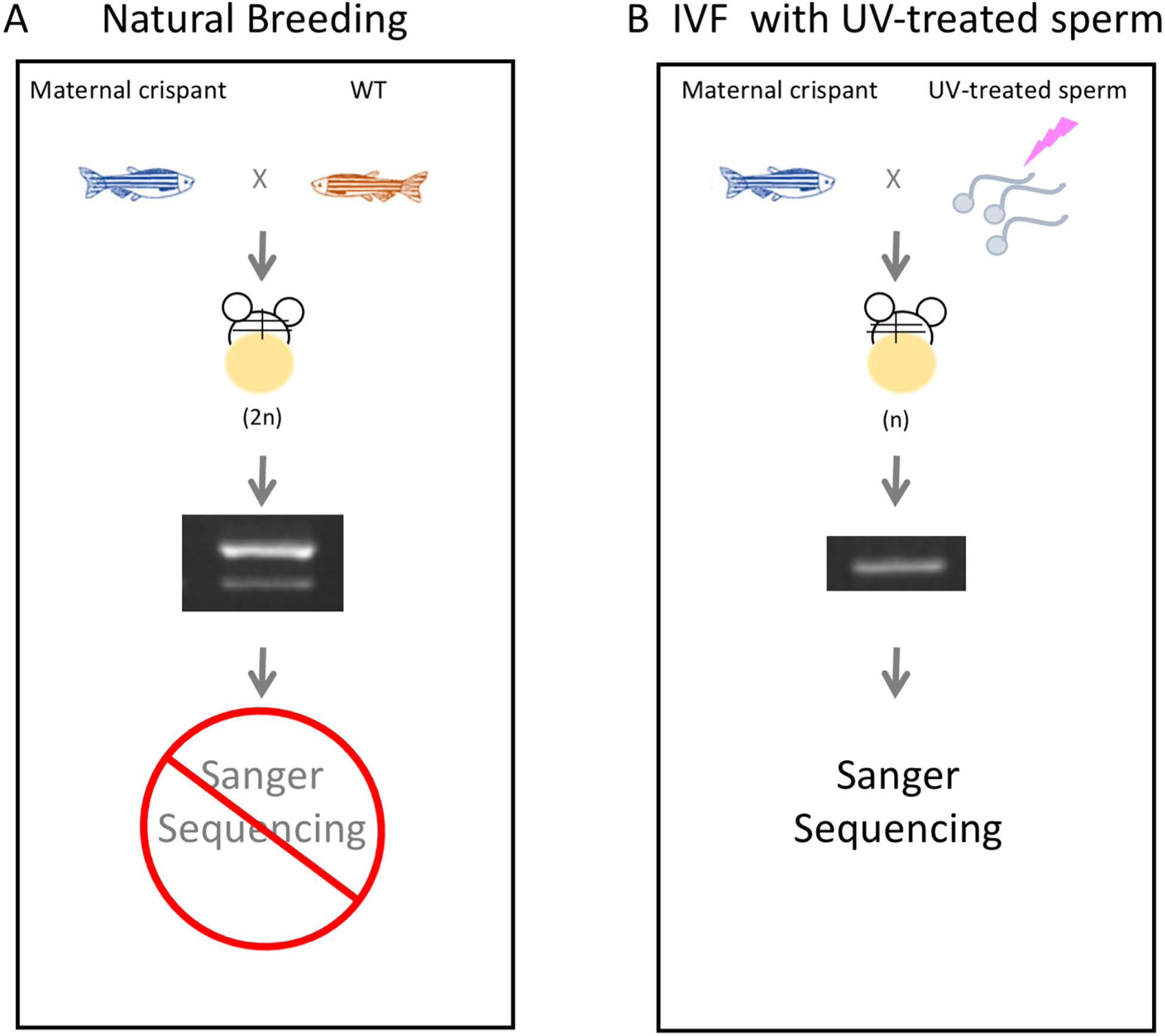
Identifying INDELS in maternal crispants. A) Embryos generated by natural breeding contain two different alleles, one from each parent. The presence of two alleles in the embryos makes it challenging to identify the genetic lesions in maternal crispants by Sanger sequencing. B) Haploid production through IVF of crispant oocytes with UV-treated sperm generates embryos with a single maternal allele, which facilitates Sanger Sequencing and genetic analysis.

For four out of the five targeted genes: *birc5b, prc1l, kpna7,* and *fhdc3,* we sequenced single maternal crispant haploid embryos from every female that presented the maternal crispant phenotype (three to thirteen embryos per female). Every sequenced maternal crispant haploid contained genetic lesions in the expected targeted gene, consistent with the germ line containing biallelic mutations in those targets (Figure 8–9; Table 3).

**Figure 8:**
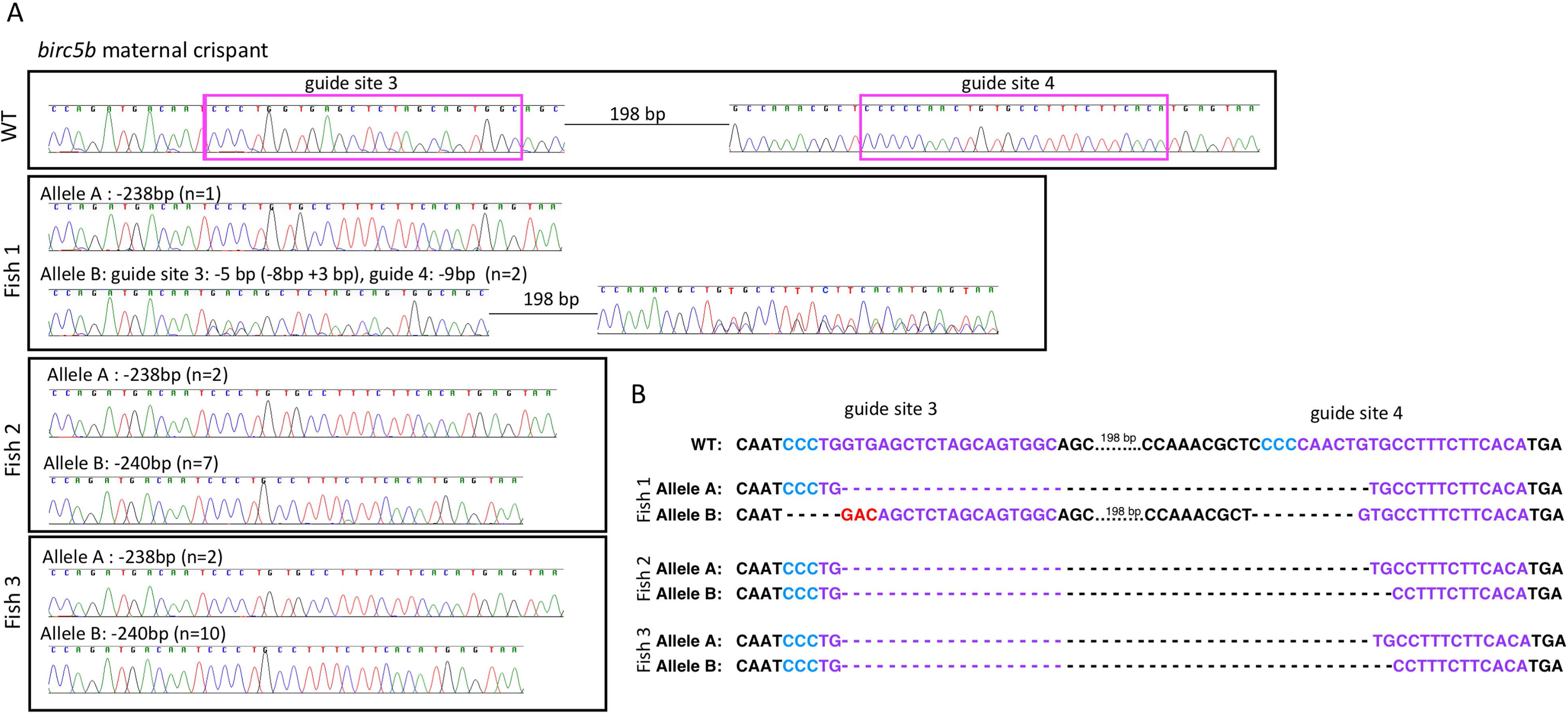
Sequencing data from haploid *birc5b* maternal crispants. A) Chromatograms of sequences for guide sites that contained INDELS after CRISPR-Cas9 mutagenesis in wild-type embryo and *birc5b* maternal crispant haploids from three different F0 fish. B) Sequence alignment of sequences corresponding to the chromatograms, with the guide site in purple and the PAM site in blue. Red text indicates the addition of new base pairs and dashes deleted bases.

**Figure 9:**
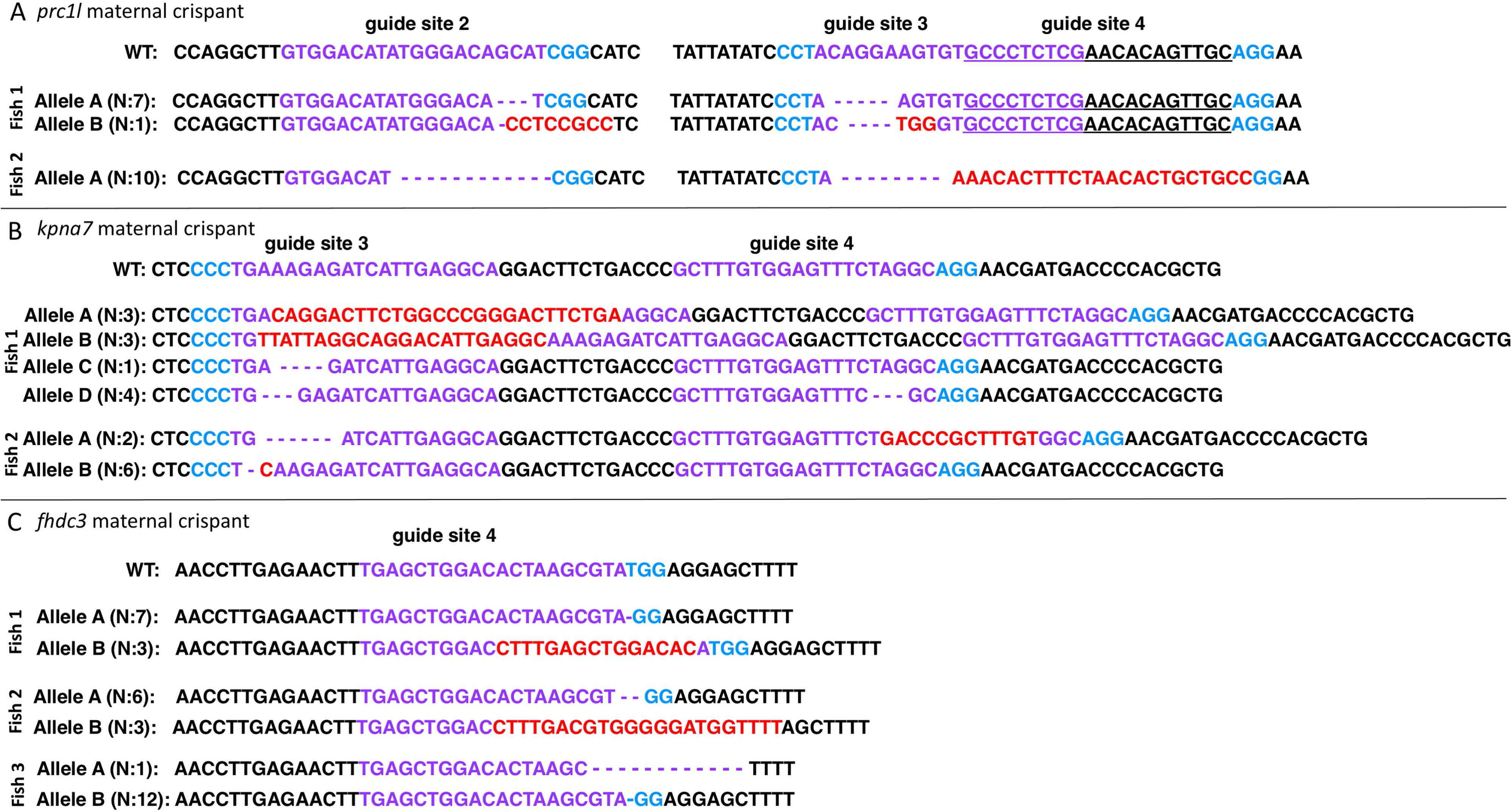
Sequencing data from maternal crispant haploids for *prc1l* (A), *kpna7* (B), and *fhdc3* (C) showing the generated lesions. Text coding as in Figure 8.

**Table 3:** Percent induction, location, and type of INDELS found in 4 of the maternal crispants.

|  | *birc5b* | *prc1l* | *kpna7* | *fhdc3* |
| --- | --- | --- | --- | --- |
| Total number of embryos sequenced | 24 | 18 | 19 | 32 |
| Total number of embryos with INDELS | 24 | 18 | 19 | 32 |
| Mutation in one site | 0 | 0 | 13 | 32 |
| Mutations in multiple sites | 24 | 18 | 6 | 0 |
| Location of INDELS |  |  |  |  |
| Guide Site 1 | 0 | 0 | 0 | 0 |
| Guide Site 2 | 0 | 18 | 0 | 0 |
| Guide Site 3 | 24 | 18 | 19 | 0 |
| Guide Site 4 | 24 | 10 | 6 | 32 |
| Types of INDELS |  |  |  |  |
| In-Frame mutation | 6 | 17 | 6 | 1 |
| Frame shift mutation | 19 | 19 | 19 | 31 |
| Average protein length | WT: 144 a.a.<br>Maternal Crispant: 95 a.a | WT: 508 a.a.<br>Maternal Crispant: 189 a.a | WT: 520 a.a.<br>Maternal Crispant: 127 a.a. | WT: 806 a.a.<br>Maternal Crispant: 131 a.a. |

In traditional zygotic crispants, it has been reported that the injected embryos are mosaic and contain multiple alleles, a majority of which are frameshifts leading to premature stop codons (Trubiroha et al., 2018; Wu et al., 2018). In these zygotic crispants, when multiple guide RNAs are used, mutations are observed in all four of the guide sites at varying frequencies, with some lesions being larger deletions spanning multiple guide sites (Wu et al., 2018). Similar to traditional crispants, edited lesions in maternal crispants include mutations in multiple guide sites as well as site spanning deletions, with a majority of DNA lesions containing frameshifts and premature stop codons (Figure 8–9; Table 3). We find that maternal crispant embryos exhibit lesions in only a subset of the guide sites (Table 3), with the same guide sites being edited in different F0 females (Figure 8–9). For example, *birc5b* maternal crispants from three different F0 females contained mutations only in guide sites three and four, and *fhdc3* maternal crispants from three different F0 females only in guide site four (Figure 8–9; Table 3). This is consistent with previous studies that show selective site usage and repair are influenced by cellular environment (Allen et al., 2018).

DNA lesions observed in single haploid maternal crispants represent a mixture of induced alleles, potentially including unique lesions from different germ cell precursors in the mosaic F0 female germ line. Additionally, the phenotype of maternal crispant embryos results from the loss of function of the two maternal alleles in the developing oocyte. Thus, sampling of targeted lesions in maternal crispants is expected to reveal multiple lesions, reflecting both the mosaicism and diploid content of the maternal germline. Consistent with this expected complexity, we observed up to four different alleles in clutches of maternal crispant embryos from single F0 females, with an average of approximately two alleles per clutch.

Assuming random segregation, each of the two maternal alleles in germ line cells containing biallelic lesions is expected to be transmitted to maternal crispants in an approximately 1:1 ratio. Interestingly, lesions identified in single maternal crispants haploids are not recovered in such equal ratios, for example all ten haploid maternal crispant embryos sequenced from *prc1l* crispant fish 2 have an identical allele (Figure 9A). This unexpected uneven recovery of germ line lesions could reflect a biased repair mechanism of CRISPR-Cas9-induced lesions in the germline, as has been observed in other systems (Allen et al., 2018; Brunner et al., 2019).

In sum, we identified genetic lesions in targeted maternal genes in all analyzed haploid embryos from F0 females, consistent with maternal crispant progeny embryos being derived from germ cells with biallelic lesions in those gene targets.

### Establishment of a stable line to confirm a maternal crispant phenotype

Confirmation that observed maternal crispants phenotypes correspond to the targeted genes requires the analysis of the phenotype in stable CRISPR-Cas9 mutant lines, an approach that we carried out for *kpna7*. We outcrossed an F0 male sibling of an F0 female that yielded maternal crispant embryos, to propagate CRISPR-Cas9 edited alleles. One such allele, *kpna7* ^uw107^, contains two mutations in the *kpna7* gene, one in guide site three (6 bp deletion) and one in guide site four (11bp deletion)(Supplementary Figure 2A), with the second mutation creating a premature stop codon (Supplementary Figure 2B). As expected, homozygous mutant females for *kpna7* ^uw107^ led to a 100% phenotype in progeny embryos similar to that of *kpna7* maternal crispants: most embryos stall at the sphere stage (Supplementary Figure 2C), with only a fraction of the embryos initiating epiboly and all embryos lysing prior to 24 hpf (n=365 embryos). The phenotype of embryos from non-mosaic *kpna7* ^uw107^ homozygous females confirms the target in the corresponding maternal crispants and provides proof-of-principle for the use of maternal crispants to identify maternal-effect genes.

## Discussion

In this study, we developed a reverse genetic approach to identify, in a single generation, novel maternal-effect genes required for early embryonic development of the zebrafish. The innovation of CRISPR-Cas9 allows targeting potential maternal-effect genes, but visualizing maternal-effect phenotypes using a standard method involving stable lines takes approximately 3 generations from the time of injection. The maternal crispant strategy described in this paper utilizes CRISPR-Cas9 to identify maternal-effect genes in a single generation, using fewer resources than a traditional CRISPR-Cas9 targeting scheme. This screening strategy is based on efficient biallelic editing abilities of CRISPR-Cas9 through multiplexing of guide RNAs (Burger et al., 2016; Jao et al., 2013; Shah et al., 2015; Shankaran et al., 2017; Trubiroha et al., 2018; Wu et al., 2018). To assay the feasibility of a maternal crispant approach, we targeted three known maternal-effect genes, *motley/birc5b* (Nair et al., 2013), *tmi*/*prc1l* (Nair et al., Submitted), and *aura*/*mid1ip1l* (Eno et al., 2016), and were able to recover at high efficiency maternal crispant embryos that showed the known maternal-effect phenotypes. Using bioinformatic selection criteria, we selected four novel candidate maternal-effect genes to target using the maternal crispant approach. Of these, targeting of *kpna7* and *fhdc3* produced unique maternal-effect phenotypes. A combination of maternal haploids and Sanger sequencing allowed us to identify and characterize INDELS in maternal crispants, facilitating the association of observable phenotypes to the gene targets. We additionally validated the *kpna7* maternal crispant phenotype using an established stable line. Our studies show that the production of maternal crispants is an effective strategy to identify maternal-effect genes.

### A maternal crispant approach allows the direct generation of maternal-effect phenotypes

Using a maternal crispant approach, we targeted three genes previously known to cause a maternal-effect phenotype when mutated, as well as four candidate genes predicted to have a maternal-effect by virtue of their expression pattern and duplicate nature. Using this approach, we were able to phenocopy all three previously known genes and identified novel phenotypes in two of the four candidates. In experiments targeting the five genes for which maternal crispant phenotypes were observed, most (14/15) screened F0 females produced embryos that showed maternal-effect phenotypes. These results suggest efficient biallelic editing of the germline for multiple targets, thus allowing for the identification of novel maternal-effect phenotypes in the progeny of injected (F0) fish.

Clutches that contained maternal crispant embryos were observed at rates that ranged from 12.5 % to 100 %, with an average rate of approximately 69 %. These clutches additionally contain phenotypically wild-type embryos, derived from oocytes without biallelic genetic edits in a mosaic F0 germ line, which can act as an internal control for variables such as fertilization time and developmental rate. The effective production of maternal crispant embryos indicates the method is successful in creating biallelic editing in the germline. Another advantage of the maternal crispant screen is that it is straightforward to simultaneously establish stable mutant lines, either from F0 sibling males, as demonstrated in the case of *kpna7* ^uw107^ allele, or from F1 offspring carrying induced lesions. These stable lines can be used to confirm phenotypes observed in the maternal crispants.

In these studies, we only identified novel maternal-effect phenotypes via gross morphological defects and this approach may allow identifying a large fraction of maternal-effect mutations. However, lethality might not be a necessary outcome for a maternal-effect phenotype. For example, the depletion of Rest/NRSF in zebrafish embryos causes the upregulation of target genes leading to changes in primary motor neuron architecture and behavior (Moravec et al., 2016). The use of transgenic lines that highlight specific structures or cell types, as has been done for heart and thyroid development in zygotic crispants (Trubiroha et al., 2018; Wu et al., 2018), would further increase the power of the approach.

### Genetic variation in maternal crispants

A survey of haploid maternal crispants showed that genetic lesions tended to involve certain guide sites but not others. In some cases, multiple independently created lesions involving identical improperly repaired sequences were found in the progeny of different F0 females. This apparent guide site selectivity contributed to limited genetic variation in maternal crispants for specific genes.

Most maternal products are produced during oocyte growth when the oocyte nucleus is in the diploid state (Reviewed in (Pelegri, 2003)). Thus, two alleles in a biallelically edited germline would be expected to be recovered in an equal ratio in haploid maternal crispants. Surprisingly, we find that a majority of edited alleles are not recovered equally in maternal crispants of the same clutch. For example, in the case of *prc1l*, ratios of recovery of two different alleles in haploid embryos were 7:1 (female 1) and 10:0 (female 2). This uneven allele recovery of alleles could be caused by a repair confirmation bias for specific INDELS (Allen et al., 2018), resulting in independent improperly repaired double strand breaks with identical sequences. A second possibility is homologous recombination/homology-directed repair, where the double stranded break in one chromatid is corrected using the other edited sister chromatid as a template (Brunner et al., 2019), creating identical mutations in both copies of the gene. We cannot rule out an alternative explanation where the observed uneven allele recovery is caused by biased segregation during meiosis, possibly caused by damage to one chromatid.

### Selection of maternal-effect candidate genes using bioinformatic approaches

We identified genes that are likely candidates to have a maternal-effect function using a combination of an RNA sequencing database (White et al., 2017) and phylogenetic analysis. In eukaryotes, whole gene duplications have allowed for significant evolutionary events to occur because the functions of the duplicated genes are less constrained (Long et al., 2003; Lynch and Conery, 2000; Ohno, 2013). In these cases, one of the gene copies can provide an essential basic function, allowing a second gene copy to acquire new functions. The zebrafish genome has undergone multiple rounds of whole-genome duplications (Meyer and Schartl, 1999), with a portion of duplicated genes experiencing functional divergence to develop new roles, including a maternal-specific function. Our studies are consistent with this phenomenon and suggest that a combination of expression data coupled to phylogenetic analysis allows for the effective identification of genes candidates with maternal-specific functions.

### Identification of new maternal-effect genes in early development

Using the above criteria, we selected 4 candidate maternal genes and tested their function using a maternal crispant approach. Out of these 4 genes, we identified new maternal-effect phenotypes in two of the genes, *kpna7* and *fhdc3*.

Past research in mice has shown that loss of function conditions for *kpna7,* an Importin α involved in nuclear transport, results in imbalanced sex ratios, with mutant females producing embryos with an embryonic-lethal phenotype (Hu et al., 2010). Knockdown of this maternally expressed transcript in other mammalian embryos, i.e., cattle and pigs, also showed reduced cellular cleavages leading to arrested embryonic development (Tejomurtula et al., 2009; Wang et al., 2012). In humans, increased expression of *kpna7* has been noted as promoting malignant properties in pancreatic cancer cell lines by regulating the formation of the mitotic spindle and nuclear morphology (Laurila et al., 2014; Vuorinen et al., 2018). We find that, in zebrafish, knocking out *kpna7* causes a 100% maternal-effect lethal phenotype involving a developmental stall starting at the sphere stage, a phenotype reminiscent of that caused by *kpna7* loss of function in mammalian species (Tejomurtula et al., 2009; Wang et al., 2012).

In zebrafish, the developmental stall caused by *kpna*7 loss function is likely due at least partly to the elimination of nuclei from a subset of cells during development, which precludes zygotic genome activation and epibolic movements in those cells (Kane et al., 1996; Knaut et al., 2000; Lee et al., 2013; Lindeman and Pelegri, 2012). Indeed, we observed abnormal DNA segregation events in early *kpna7* maternal crispant early embryos, likely responsible for generating cells that lack DNA at later stages. Kpna7 has been found to be required for nuclear lamin import and nuclear lamina organization (Vuorinen et al., 2018), and it is possible that nuclear envelope reassembly, which in the rapid cell cycles of the early zebrafish embryo occurs concurrently with DNA segregation (Abrams et al., 2012; Lindeman and Pelegri, 2012), generates a link between *kpna7* function in nuclear import and DNA segregation.

A second maternal crispant target in this study was *fhdc3*, a maternally expressed homolog of the inverted formin *fhdc1*. *fhdc3* maternal crispants exhibit an exaggerated constriction of the cytokinetic ring. In wild-type, this actin-based ring forms at the margin of the blastodisc during the early cleavage stages and is thought to drive ooplasmic flow and yolk segregation during cycles of contraction and relaxation associated with the cell cycle (Fuentes and Fernández, 2010; Leung et al., 2000). The exaggerated constriction phenotype in *fhdc3* maternal crispants is no longer observed after the 6^th^ cell cycle, a time coincident with the disappearance of the cytokinetic ring in wild type. However, *fhdc3* crispant embryos continue to exhibit defects as reflected by ectopic yolk inclusions in the embryo proper and reduced embryonic viability. Thus, *fhdc3* is a maternal gene with a temporarily restricted yet essential function in the regulation of the actin-based cytokinetic ring during the earliest embryonic cell cycles, which contributes to proper segregation of yolk from cytoplasm.

We did not identify maternal-effect phenotypes for two targeted genes, the *spire-*family gene *zgc:114123* and the S subfamily potassium channel *kcns3b,* in spite of efficient production of INDELS in somatic cells of injected F0 embryos. Considering the expression pattern of *zgc:114123* and role of Spire in oogenesis in other systems (Bradley et al., 2020; Leader et al., 2002; Manseau and Schüpbach, 1989; Pfender et al., 2011), it is possible that a loss of function for *zgc:114123* affects oocyte development, precluding obtaining maternal crispant embryos in F1 clutches. The number of maternal transcripts for *kcns3b* is significantly less than other targets in our studies, so that the absence of maternal crispants for this gene may reflect a minor or non-essential role for this gene during development.

In summary, this study presents an effective approach to produce embryos with loss of function for maternal-effect genes, taking advantage of a high rate of CRISPR-Cas9-induced biallelic editing in the germline. The resulting maternal crispants allow for the identification and characterization of maternal-effect genes in a single generation. In teleost fish, a subset of maternal genes has arisen from gene duplication events, and targeting maternal-specific duplicate genes could provide insight into the role of gene homologs in development and evolution. Thus, a maternal crispants approach can help expedite our understanding of the role of maternal products during development.

## Acknowledgments

We thank past and current members of the Pelegri lab for their contributions to our research, particularly our animal husbandry staff for the care of the aquatic facility. We are also grateful for the comments and insight on the manuscript by Christina Hansen and Ryan Trevena. Microscopy was performed at the Newcomb Imaging Center, Department of Botany at the University of Wisconsin-Madison. Funding was provided by NIH grant to F.P. (GM065303)

---

**Movie 1:** Cellular cortex of a *kpna7* maternal crispant at 6 hpf. Individual focal planes of confocal Z-stack of a 6hpf *kpna7* maternal crispant embryo (shown as a 2D Z-projection in Figure 4B.) The cell membranes are labeled with anti-β-catenin antibodies (red) and DNA is stained with DAPI (blue). Visualization of sequential focal planes demonstrates that the *kpna7* maternal crispants contain cells that lack nuclei.

**Movie 2:** Cellular cortex of a wild-type embryo at 6 hpf. Individual focal planes of confocal Z-stack of 6hpf wild-type embryo (shown as a 2D Z-projection in figure 4B.) The cell membranes are labeled with anti-β-catenin antibodies (red) and DNA is stained with DAPI (blue).

**Supplementary Figure 1:** mRNA expression levels and gene structures for candidate maternal-effect genes, *zgc114123* and *kcn3b*, which did not display a phenotype in the maternal crispants. A, C) Expression pattern for the *spire* (A) and *kcn3*(C) families during early development, from zygote to larval day 5. For each family, the maternal-specific transcripts are represented in red and the gray bar marks zygotic genome activation. (B, D) Gene structure diagrams of *zgc114123* (B) and *kcn3b* (D), with guide RNA target sites (red lines) and PAM sites (red stars).

**Supplementary Figure 2**: Validation of the *kpna7* maternal crispant phenotype using a stable Crispr-Cas9 line. A) Chromatograms and sequencing alignments for wild-type and *kpna7* ^uw107/UW107^ samples lesions in edited guide sites. In the sequencing alignment, the purple text represents the guide sites, and the blue text is the PAM site. B) Diagrams highlighting known conserved domains in the Kpna7 protein and the predicted truncated product encoded by the *kpna7 ^UW107^* allele and amino acid sequences showing the predicted changes and appearance of a premature stop codon. C) Embryos from *kpna7 ^UW107/UW107^* females exhibit a fully penetrant phenotype identical to that observed in *kpna7* maternal crispants.

**Supplementary Table 1:** Candidate genes *zgc:114123* and *kcn3b* did not result in any apparent maternal crispant embryos, observed after Sanger sequencing of maternal crispant and wild type haploid embryos.

**Excel file 1:** Oligos for guide RNAs and primers used in this study.

## Notes

### Competing Interest Statement

The authors have declared no competing interest.

